# Reconstructing the Transcriptional Regulatory Network of Probiotic *L. reuteri* is Enabled by Transcriptomics and Machine Learning

**DOI:** 10.1101/2023.07.03.547516

**Authors:** Jonathan Josephs-Spaulding, Akanksha Rajput, Ying Hefner, Richard Szubin, Archana Balasubramanian, Gaoyuan Li, Daniel C. Zielinski, Leonie Jahn, Morten Sommer, Patrick Phaneuf, Bernhard O. Palsson

## Abstract

I

*Limosilactobacillus reuteri*, a probiotic microbe instrumental to human health and sustainable food production, adapts to diverse environmental shifts via dynamic gene expression. We applied independent component analysis to 117 high-quality RNA-seq datasets to decode its transcriptional regulatory network (TRN), identifying 35 distinct signals that modulate specific gene sets. This study uncovers the fundamental properties of *L. reuteri’s* TRN, deepens our understanding of its arginine metabolism, and the co-regulation of riboflavin metabolism and fatty acid biosynthesis. It also sheds light on conditions that regulate genes within a specific biosynthetic gene cluster and the role of isoprenoid biosynthesis in *L. reuteri’s* adaptive response to environmental changes. Through the integration of transcriptomics and machine learning, we provide a systems-level understanding of *L. reuteri’s* response mechanism to environmental fluctuations, thus setting the stage for modeling the probiotic transcriptome for applications in microbial food production.

**Graphical Abstract:** 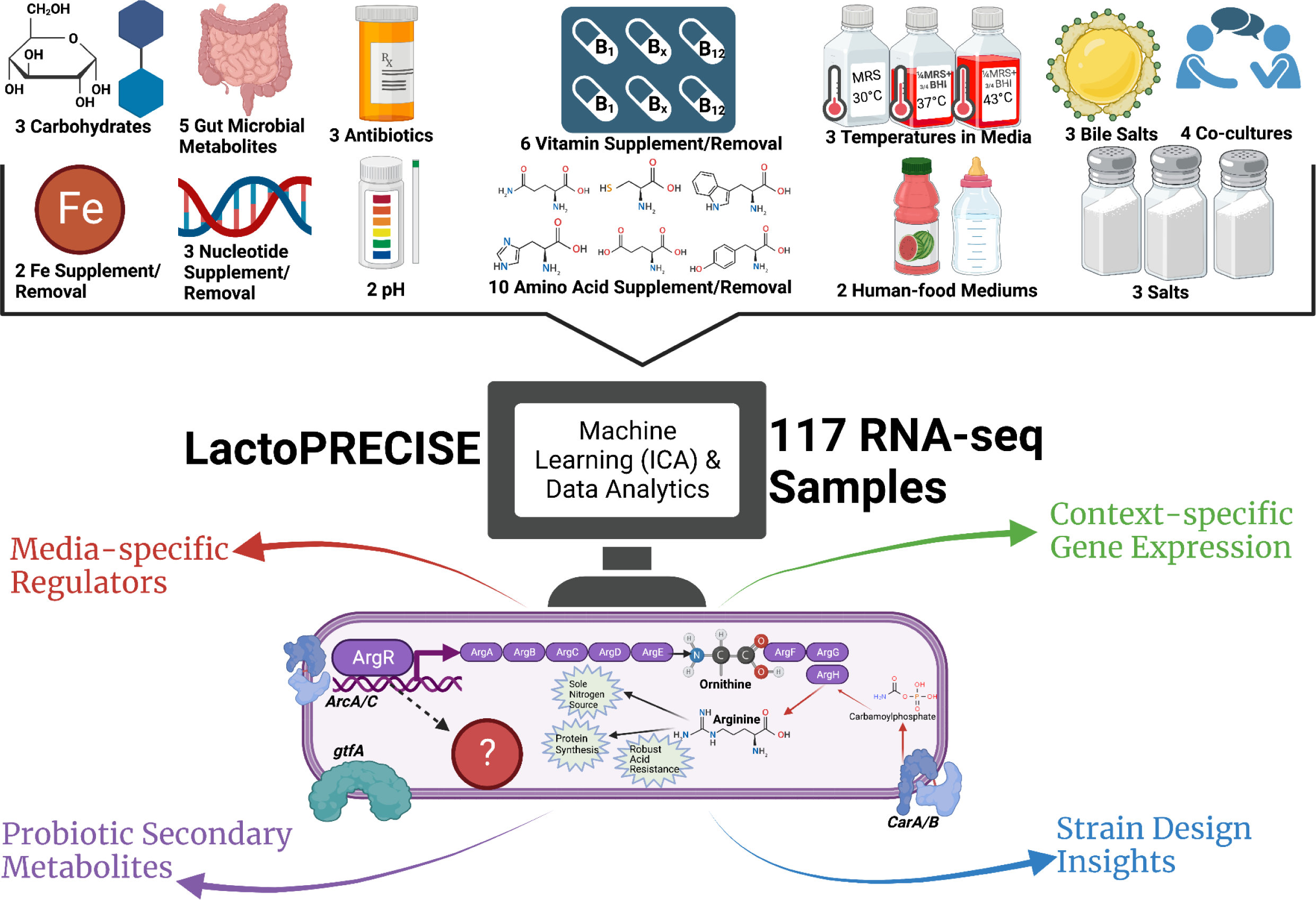

Comprehensive iModulon Workflow Overview. Our innovative workflow is grounded in the analysis of the LactoPRECISE compendium, a curated dataset containing 117 internally sequenced RNA-seq samples derived from a diversity of 50 unique conditions, encompassing an extensive range of 13 distinct condition types. We employ the power of Independent Component Analysis (ICA), a cutting-edge machine learning algorithm, to discern the underlying structure of iModulons within this wealth of data. In the subsequent stage of our workflow, the discovered iModulons undergo detailed scrutiny to uncover media-specific regulatory mechanisms governing metabolism, illuminate the context-dependent intricacies of gene expression, and predict pathways leading to the biosynthesis of probiotic secondary metabolites. Our workflow offers an invaluable and innovative lens through which to view probiotic strain design while simultaneously highlighting transformative approaches to data analytics in the field.

## II. Introduction

*Limosilactobacillus reuteri (L. reuteri*) has emerged as a subject of intensive investigation, primarily owing to its multifaceted roles in commercial applications and human health. It is extensively harnessed in a variety of consumer goods as a fermentation agent, producing the natural antimicrobial reuterin that efficiently thwarts food-borne pathogens such as yeasts, molds, and a diverse array of bacteria, thereby preventing food spoilage (1, 2). Moreover, *L. reuteri* contributes to the improvement of food taste via the production of certain fatty acids like linoleic acid and enriches the nutrient content and flavor profile of food products through the synthesis of vital vitamins or amino acids (3, 4).

In addition, there is growing recognition that *L. reuteri* is a crucial constituent of the human-associated microbiome, where it displays its prowess by mitigating the presence of harmful microbes and fostering beneficial commensals (2,5). *L. reuteri* can be found in the human gut, vagina, and breast milk, where it provides a plethora of health benefits pertaining to host-associated microbiome modulation (6). Specifically, the benefits of *L. reuteri* range from improving vaginosis symptoms (7), reducing serum fat/cholesterol levels (8, 9), alleviating chronic intestinal inflammation (10), ameliorating symptoms of cystic fibrosis (11), protecting the gut microbiome of extremely preterm infants (12), and even alleviating maternal depression (13), among a host of other advantages (6).

However, for a significant proportion of these observed beneficial effects attributed to *L. reuteri*, the underlying mechanisms of action remain largely unexplored. Generally, the health impact of this organism has been intricately linked to metabolic activity, which is, in turn, influenced by environmental conditions. Consequently, to establish a more robust understanding of how environmental conditions affect the adaptation of *L. reuteri* gene expression, we have employed the machine learning algorithm independent component analysis (ICA) on a compendium of *L. reuteri* gene expression data.

Machine learning, particularly ICA, has the capability to integrate a wide array of data inputs by extracting focal points of interest (14). In this study, we applied ICA to elucidate the activity of iModulons (15), representing groups of genes that provide a comprehensive view of broad biological functions in relation to varying experimental conditions. Notably, the transcriptional regulatory network (TRN) of a given organism holds the potential to perceive intricate environmental stimuli, coordinating cellular gene expression in a bottom-up manner (16). Conversely, reverse engineering the TRN can shed light on the organism’s adaptive response to a dynamic environment (17, 18, 19). Therefore, the reconstruction of *L. reuteri’s* TRN is critical, not only for prediction but also for gaining mechanistic insights into its adaptive response to various environments, from industrial settings to the human microbiome.

Despite the widespread utilization of *L. reuteri* for health and commercial benefits, there exists a notable gap in studies attempting to elucidate the metabolic mechanisms underpinning this organism’s response to dynamic conditions. As of this writing, while investigating NCBI’s SRA database, it was found that only 36 studies have explored *L. reuteri* gene expression in response to changing environmental conditions (**Figure 1D**). Of these, only six leveraged next-generation RNA sequencing, and just one utilized RNA-seq to study *L. reuteri in vitro*. To bridge this gap, we have created the LactoPRECISE compendium, based on high-quality RNA-seq datasets spanning a wide array of media conditions, in order to reconstruct a genome-scale TRN. This compendium, consisting of 117 individual experiments, enables us to interrogate gene expression regulators and characterize their activity. Fundamentally, quantitative measurements of expression profiles reflect a broad amalgamation of the activators of a specific transcriptional regulator under a given condition. This information, coupled with ICA, can shed light on the context-specific TRN structure and establish key differences from context-specific gene expression at a genome scale. Specifically, only 45% of the gene expression variance from our compendium can be explained, indicating that ICA results should be interpreted cautiously in the context of biological phenomena, especially in the absence of robust gene-level annotations.

**Figure 1:**
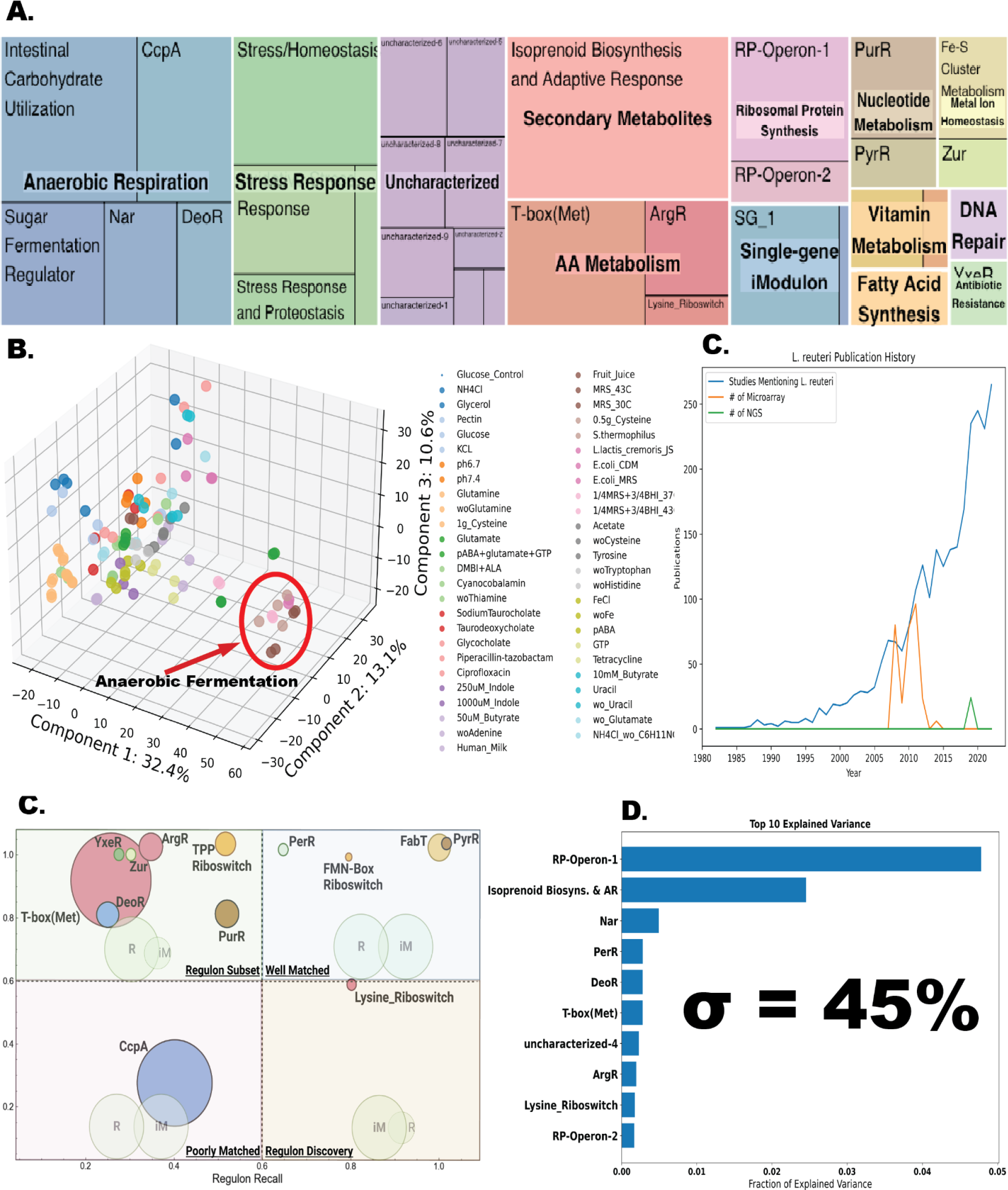
Description of the LactoPRECISE transcriptomics Compendium. A) A treemap depiction of the 35 identified iModulons and their associated functional categories. The dimensions of the individual boxes correlate with the number of genes found in each respective iModulon. B) A PCA plot based on the compendium gene expression highlights conditions that are similar or different from one another. C) A comprehensive scatterplot of all Regulatory iModulons. Each point signifies an individual iModulon, with the size of the point proportionate to the gene count within the corresponding iModulon. Regulatory iModulons are segmented into four quadrants: regulon subset (top left), well-matched (top right), poorly matched (bottom left), and regulon discovery (bottom right). High iModulon recall and regulation recall values denote a robust consistency between a particular iModulon and a previously characterized one. The overlap between the gene lists in computed iModulons and regulons within each quadrant is intuitively visualized by background Venn diagrams in each quadrant. Regulon recall can be defined as the ratio of shared genes between an iModulon and a regulon to the total genes in a regulon, whereas iModulon recall is the proportion of shared genes to the overall genes in an iModulon. D) Graphical representation of the volume of published research that includes the mention of ‘*L. reuteri’* in the abstract, plotted against the body of published microarray or next-gen RNA-seq literature. E) Highlight of the top 10 iModulons contributing the greatest proportion of explained variance. These iModulons encapsulate key functions such as ribosome protein synthesis, secondary metabolite biosynthesis, stress response, amino acid synthesis, fatty acid synthesis, and unknown functions.

Despite these limitations, the application of ICA to RNA-seq data has shown promise in unearthing novel biological insights into *L. reuteri*. In this study, we successfully applied ICA to the LactoPRECISE compendium to define iModulons, which greatly improve our understanding of *L. reuteri* transcriptional regulation; more effort is still required to completely reconstruct the TRN of this organism. This analysis has identified a collection of Functional iModulons that describe biological functions across various conditions. Importantly, many of these iModulons can be linked to the regulatory control of microbial food production, providing valuable insights into the regulatory mechanisms and conditions controlling the production of specific value-added probiotic products. Specifically, we have provided insight into the regulation of *L. reuteri* modules that are related to microbial food production, which are characterized by the iModulons: *ArgR* (20, 21), *CcpA* (22), *DeoR* (23), *FabT* (24), FMN-Box Riboswitch (25), Isoprenoid Biosynthesis & Adaptive Response (26), Lysine Biosynthesis (27), *PyrR* (28), T-Box (Methionine) (29), and TPP Riboswitch (30). At least two of these iModulons have demonstrated potential co-regulation between modules, highlighting shared functions that may prove useful for future strain design initiatives to control beneficial fatty acid and vitamin production (31). This study also sought to uncover the TRN from a gram-negative bacteria, these results were compared with those from gram-positive microbes to identify core similarities and differences within similar iModulons through the utilization Bitomics (32). Finally, iModulons have offered data-driven insights into the conditions that activate or repress specific genes captured within iModulons. This result demonstrates the utility of data-driven, systems-level experimentation in informing and engineering strain design by controlling gene regulation (33, 34).

## III. Methods

### A. RNA Extraction & Library Preparation

In this study, we used *Limosilactobacillus reuteri* MM4-1A (PTA-6475) as the target strain. The strain was cultured in different growth media, including the base medium CDM, MRS, and specific food-like media, such as human milk, infant formula, or melon fruit juice, with appropriate supplements. We conducted experiments under aerobic or anaerobic conditions, covering a total of 50 unique conditions, which included variations in carbohydrate supplementation, gut microbial metabolites, salt stress, vitamin supplementation or removal, various media compositions, temperature changes, bile salt stress, co-cultures with other microbes, iron supplementation or removal, nucleotide supplementation or removal, pH changes, amino acid supplementation or removal, human-food media, and antibiotic treatments. A detailed list of all the growth conditions can be found in **Supplementary Table 1.**

To prepare the samples, overnight cultures of the strains were grown at defined temperatures with mixing in the respective media. The cultures were then diluted to a starting optical density at 600 nm (OD600) of approximately 0.01 and incubated at defined temperatures with stirring. When the cultures reached an OD600 of 0.4, 2 ml were transferred to centrifuge tubes containing 4 mL of RNAprotect Bacteria Reagent (Qiagen). After vortexing for 5 seconds, the tubes were incubated at room temperature for 5 minutes. Subsequently, the samples were centrifuged for 10 minutes at 5000 × g, and the supernatant was removed. The cell pellets were stored at −80°C until further processing. In conditions involving antibiotic treatment, the antibiotics were added at 2× or 5× their minimum inhibitory concentration (MIC) when the bacterial culture reached an OD600 of approximately 0.2. The cultures were then incubated at defined temperatures with stirring for an additional hour before sample collection.

Total RNA was isolated and purified from the frozen cell pellets using the Zymo Research Quick-RNA Fungal/Bacterial Microprep Kit, following the manufacturer’s protocols. To remove ribosomal RNA (rRNA), 1 μg of total RNA was treated with thermostable RNase H (Hybridase) and short DNA oligos that specifically targeted and degraded rRNA at 65°C to preserve mRNA integrity. The resulting rRNA-subtracted RNA was converted into libraries using the KAPA RNA HyperPrep kit, which incorporated short Y-adapters and barcoded PCR primers. The libraries were quantified using a fluorescent assay and assessed for size distribution using a TapeStation. After pooling the libraries and performing a 1× SPRI bead cleanup to remove residual PCR primers, the final library pool was quantified and sequenced on an Illumina instrument (NextSeq, Novaseq).

### B. The Formation and Composition of LactoPRECISE

Given the scarcity of comprehensive in vitro studies on *L. reuteri* using solely next-generation RNA-seq in public repositories, the data for LactoPRECISE was exclusively sourced from internal sequencing. The RNA-seq samples generated in this study were prepared following the PyModulon workflow delineated by Sastry et al. 2021 (35). The replication of this pipeline is made possible through the access code at https://github.com/avastry/modulome-workflow. As the final product of this process encompasses all currently computable iModulons from this organism, we have defined the resultant database as “LactoPRECISE.”

### C. RNA-Seq Data Acquisition, Processing, and Management

In brief, the 163 raw paired-end RNA-seq samples generated for this study were subjected to the prokaryotic RNA-seq processing pipeline outlined by Sastry et al. 2021 (available at https://github.com/avastry/modulome-workflow) and mapped to our reference genome (GCF_020785475.1) (35). Post-processing, the gene expression compendium was log-transformed to Transcripts per Million (logTPM). Adhering to Sastry et al. 2021 quality control protocol, we manually curated the experimental metadata to distinguish biological replicate samples and annotate individual samples with details such as media descriptions, treatment specifics, environmental changes, growth stages, and other pertinent growth-related parameters (35).

We maintained strict quality control measures throughout our analysis. Data failing any of the four FASTQC measures—per base sequence quality, per sequence quality scores, per base n content, and adapter content—were excluded from further analysis. Samples with less than 500,000 reads mapped to coding sequences were likewise discarded. We employed hierarchical clustering to identify samples with atypical expression profiles. After these initial quality control steps, we carried out manual metadata curation on the remaining data. We extracted details such as strain-description, base media, carbon source, treatments, and temperature from existing literature. We assigned a unique, succinct name to each project, and within each project, we used unique identifiers to recognize biological and technical replicates. Post-curation, we removed samples if they (a) lacked metadata, (b) were without replicates, or (c) showed a Pearson R correlation between replicates lower than 0.80. The log-TPM data within each project was then centered on a reference condition specific to that project.

The gene expression data were processed through a correlation-based filtering and averaging procedure to manage dissimilarity within the samples of each group. The procedure was initiated by organizing gene expression data into groups corresponding to specific conditions. Each group was then assessed for internal consistency by computing the pairwise Pearson correlation coefficient for samples within the group. To address discrepancies in the correlation of gene expression data within each group, a specified correlation threshold was set. We implemented a range of threshold values (0.8, 0.85, 0.9, 0.95) (lower thresholds were deemed low replicate quality and thus not considered for inclusion) to analyze the sensitivity of the downstream results to this parameter. If a pair of samples within a group had a correlation coefficient lower than the threshold, the group was deemed to have significant internal variation. To counter this, we averaged all samples in the respective group to create a single, representative expression profile. The output of this process was a filtered expression matrix in which groups with substantial internal dissimilarity were represented by a single, averaged profile. This matrix was then used in subsequent analyses. Separate runs of the procedure were performed for each correlation threshold, allowing for the comparison of results across different stringency levels in the filtering process Overall, a threshold of R = 0.80 was selected for analyzing downstream ICA results (**Supplementary Figure 1B).** This extensive curation resulted in our final compendium comprising high-quality *L. reuteri* incorporating 117 RNA-seq samples.

Simultaneously, a preliminary TRN was constructed. This process entailed accumulating known transcription factor data from RegPrecise and collating regulatory interactions from existing *L. reuteri* literature (36). The resources utilized for this draft TRN development, including transcription factors, are cataloged in **Supplementary Table 2**. This integrative approach ensured a comprehensive and accurate representation of the RNA-seq data and the associated regulatory elements.

### D. Implementation of Independent Component Analysis

Following the PyModulon workflow, we employed ICA on the RNA-seq compendium using the optICA extension, which is derived from the FastICA algorithm. The optICA script employed in this process is available at https://github.com/avsastry/modulome-workflow/tree/main/4_optICA. This script generates two matrices, M and A, which are crucial in assessing iModulon activity. The M Matrix comprises robust components constituting the iModulons, while the A Matrix encapsulates the corresponding iModulon activities. Both these matrices are approximations extrapolated from the X Matrix, representing the expression data drawn from the RNA-seq compendium.

Determining the optimal number of iModulons for the compilation of LactoPRECISE required the identification of an optimal dimensionality to circumvent both under and over-decomposition of the gene expression compendium. To pinpoint this optimal dimension (37), we performed multiple rounds of gene expression profile clustering within a range of 30 to 100, using a step size 10. The optimal dimension of 60 was eventually selected, corresponding to the number of non-single gene independent components matching the final component count (**Supplementary Figure 1A**). The execution of the ICA algorithm ultimately yielded a total of 35 robust iModulons for *L. reuteri*.

### E. iModulon Regulon Enrichment, Functional Annotation, and Characterization

Regulator enrichments were determined following the methodology outlined by Sastry et al. 2021 (35). The gene annotation pipeline is available at https://github.com/SBRG/pymodulon/blob/master/docs/tutorials/creating_the_gene_table.ipy <u>nb</u>. To elucidate the biological roles of the iModulons identified within the LactoPRECISE, we utilized the PyModulon Python package (35). We first aligned each iModulon with a preliminary version of the reconstructed TRN table to discern iModulons with a substantial intersection with previously characterized transcriptional regulators. This step was critical for establishing the potential regulatory roles of the identified iModulons.

Following this, to further interpret the functional context of the iModulons and their associated genes, we KEGG (38) and Cluster of Orthologous Groups (COG) data were retrieved via the EggNOG mapper (39). UniProt IDs were secured using the Uniprot ID mapper (40), and operon information was gathered from Biocyc (41). We procured Gene Ontology (GO) annotations from AmiGO2 (42). The known TRN was accessed from RegPrecise (36) and supplemented by a manual curation of related literature. We assessed the efficacy of the predicted iModulons using ‘iModulon recall’ and ‘regulon recall’ metrics. The ‘iModulon recall’ measures the ratio of shared genes to the total genes in an iModulon, while the ‘regulon recall’ quantifies the ratio of shared genes to the total genes in a regulon (**Figure 1C**). This comprehensive characterization provides a robust framework for understanding the biological significance of iModulons within the context of *L. reuteri’s* cellular processes. Details on each iModulon and their respective functions, number of genes, and other relevant characteristics can be found in **Supplementary Table 3**.

### F. Reconstructing *L. reuteri* ArgR Bitome

Considering the lack of accessible information about gene-operon relations, we undertook an algorithmic approach to segregate genes into operons based on spatial proximity. Initially, genes were batched based on their respective accession IDs. The sorted set of genes was parsed through an iterative operation, which checked for spatial congruity with the preceding gene. If the spatial gap fell within a predefined threshold, it was assigned to the current operon; otherwise, a new operon cluster was initiated. This process ultimately led to the formation of operon clusters, which are essentially nested lists of genes grouped based on their operon classification.

To form a position-specific scoring matrix (PSSM) for ArgR, we aligned the binding sites of *E. coli’s* ArgR iModulon. We subsequently quantified the frequency of each nucleotide at individual positions, normalizing this against the background frequency in the entire sequence, which resulted in the computation of log-odds scores. The ‘*bitome_fasta.motif_search’* function was employed for the iterative computation of motif scores for equidistant segments, equivalent to the motif length. The segment with the most substantial score was earmarked as the motif score of that sequence.

Using the motif scores as the input matrix and binarized ArgR regulon memberships as target labels, a logistic regression model was built incorporating an elastic net penalty (with an L1 to L2 ratio of 0.5) for regularization. To tackle class imbalance, we employed a SMOTETomek resampling technique, set to a parameter of k_neighbors=5, to generate a balanced training set. To ensure the representation of each class, we utilized a 5-fold cross-validation with stratified sampling. The train-test-split was conducted within the operon space, ensuring genes from the same operon were kept together. The model’s performance was evaluated using an AUC-ROC curve, with a pre-decided threshold of 0.8 as the benchmark for satisfactory performance.

### G. Prediction of Biosynthetic Gene Clusters

Biosynthetic gene clusters (BGCs) in *L. reuteri* were predicted using the antiSMASH algorithm 7.0 (43). The *L. reuteri* reference genome (GCF_020785475.1) was utilized with a ‘relaxed’ detection strictness setting. AntiSMASH accurately identifies two types of *L. reuteri* BGCs, RiPP (ribosomally synthesized and post-translationally modified peptides) and Type III Polyketide Synthase (T3PKS) Additionally, the algorithm provides gene ontology annotations for the components of the predicted BGCs. Since genes within RiPP were found to be inactive in our compendium, our analysis focused on T3PKS.

### H. Generating iModulonDB Dashboards

iModulonDB dashboards were generated using the PyModulon package (35). The link to the lactoPRECISE compendium can be found here https://imodulondb.org/dataset.html?organism=l_reuteri&dataset=lactoprecise

## IV. Results

Deep Insights into the *L. reuteri* Transcriptional Regulatory Network Unveiled by Transcriptomic Compilation and Independent Component Analysis

To understand the global transcriptional regulation that governs the physiological processes of *L. reuteri*, we applied ICA to a high-quality RNA-seq compendium encompassing 117 RNA-seq samples across a range of predetermined conditions. This analysis identified 35 integral components or iModulons. Collectively, these 35 iModulons account for 45% of the variance in gene expression across our compendium and thus represent the variation derived from transcriptional regulation (**Figure 1**).

Notably, the variance explained by the 35 iModulons is significantly lower than that observed when using principal component analysis (PCA) on the **X** matrix alone. However, unlike PCA, which often lacks biological interpretation, ICA provides more nuanced insights into the biological regulation of gene expression. Through manual characterization, we can interpret the iModulons derived from independent components as sources of biologically relevant expression observable in the compendium, capturing a broad spectrum of biological functions (**Figure 1A**). S**upplementary Figure 2** illustrates the correlation of iModulons across the compendium by utilizing hierarchical clustering. **Supplementary Figure 3** further identifies the top five most significant iModulon clusters and highlights iModulons with the most similar cellular responses in *L. reuteri*.

In our initial characterization of these 35 iModulons, we built a draft TRN of *L. reuteri* using the RegPrecise database (36) and assorted literature sources that describe known *L. reuteri* transcription factors. We found that 13 of the 35 identified iModulons overlap significantly with known regulators, suggesting that these Regulatory iModulons reflect the basic structure of known regulons. Moreover, public information on TRNs can help initiate the process of reconstructing TRNs from RNA-seq data. We can further quantify the relationship between Regulatory iModulons and known regulons used to reconstruct our draft TRN by using metrics such as ‘iModulon Recall,’ which measures the fraction of genes in an iModulon that also belong to a regulon, and ‘Regulon Recall,’ the fraction of regulon genes in a specific iModulon. Many identified iModulons show a high iModulon Recall rate, indicating strong agreement between the draft TRN structure and the newly reconstructed iModulon TRN for *L. reuteri* (**Figure 1C**).

Next, we sought to characterize the remaining 22 iModulons using data from public databases such as GO, KEGG, and BioCYC (38, 41, 42). Here, we identified 11 Functional iModulons—those containing gene groups related to a specific biological function but not directly linked to a known transcriptional regulator. Two additional iModulons, dominated by a single, high coefficient gene, could be termed ‘single-gene iModulons,’ possibly regulated solely by that gene. Initially, 20 ‘uncharacterized iModulons’ were observed. These iModulons were defined through various approaches, such as investigating gene membership, use of functional gene annotations, iModulon activity against various conditions, and using ChatGPT 4.0 (45). While ICA exhibits a high success rate in identifying biologically useful modules of the TRN, many of these modules remain uncharacterized; however, utilization of ChatGPT 4.0 drastically improved our ability to functionally describe ‘Uncharacterized iModulons.’ Specifically, all genes found within the uncharacterized iModulons, in addition to their description against conditions, and an assessment of potential gene interactions with known regulons was used to identify 11 additional iModulons. Overall, nine ‘uncharacterized iModulons’ could not be categorized or adequately characterized, making them ideal targets for future research; specifically, uncharacterized-9 will be presented later in the manuscript as a potential regulator of nitrogen metabolism (**Figure 6**).

To provide a holistic view of our findings, we assigned functional categories to all iModulons (**Figure 1A**). Notably, iModulons with known biological functions that explain the highest fraction of expression variance in our compendium include those controlling ribosomal protein synthesis (RP-Operon-1), secondary metabolite production (Isoprenoid Biosynthesis and Adaptive Response), stress response (PerR), nitrate reductase (Nar), amino acid synthesis (T-box(Methionine), ArgR, Lysine Riboswitch), and anaerobic respiration (Nar, DeoR).

In **Figure 1**, we provide a comprehensive visualization of our findings. **Figure 1A** displays a treemap of the 35 identified iModulons and their functional categories, with the size of the boxes corresponding to the gene count in each iModulon. **Figure 1B** provides a PCA plot highlighting similar and distinct conditions based on gene expression in the compendium. **Figure 1C** offers a scatterplot of all Regulatory iModulons, segmented into four quadrants—regulon subset, well-matched, poorly matched, and regulon discovery—based on their level of agreement with previously characterized iModulons. **Figure 1D** represents the volume of published research on *L. reuteri* compared to the body of microarray or next-gen RNA-seq datasets, in addition to published literature. Finally, **Figure 1E** highlights the top 10 iModulons contributing to the explained variance, encapsulating functions like ribosome protein synthesis, secondary metabolite biosynthesis, stress response, amino acid synthesis, fatty acid synthesis, and unknown functions.

### A Systems-level View of *L. reuteri* iModulons Sheds Light on their Activity Across the Compendium

Given the relatively small number of iModulons in *L. reuteri* as compared to other strains (likely attributable to its small genome size) (46), it is both feasible and illuminating to delve into the specific role of each iModulon in regulating *L. reuteri* as a probiotic organism. Carefully examining individual iModulons within LactoPRECISE provides a system-level perspective on the molecular machinery that orchestrates the organism’s function. Hence, we expanded and clarified the contribution of individual genes, their weight within their respective iModulons, and the iModulon activity across all conditions for 11 iModulons related to microbial foods. This enabled us to demonstrate how iModulon results can be leveraged to generate hypotheses (**Figure 2**).

**Figure 2:**
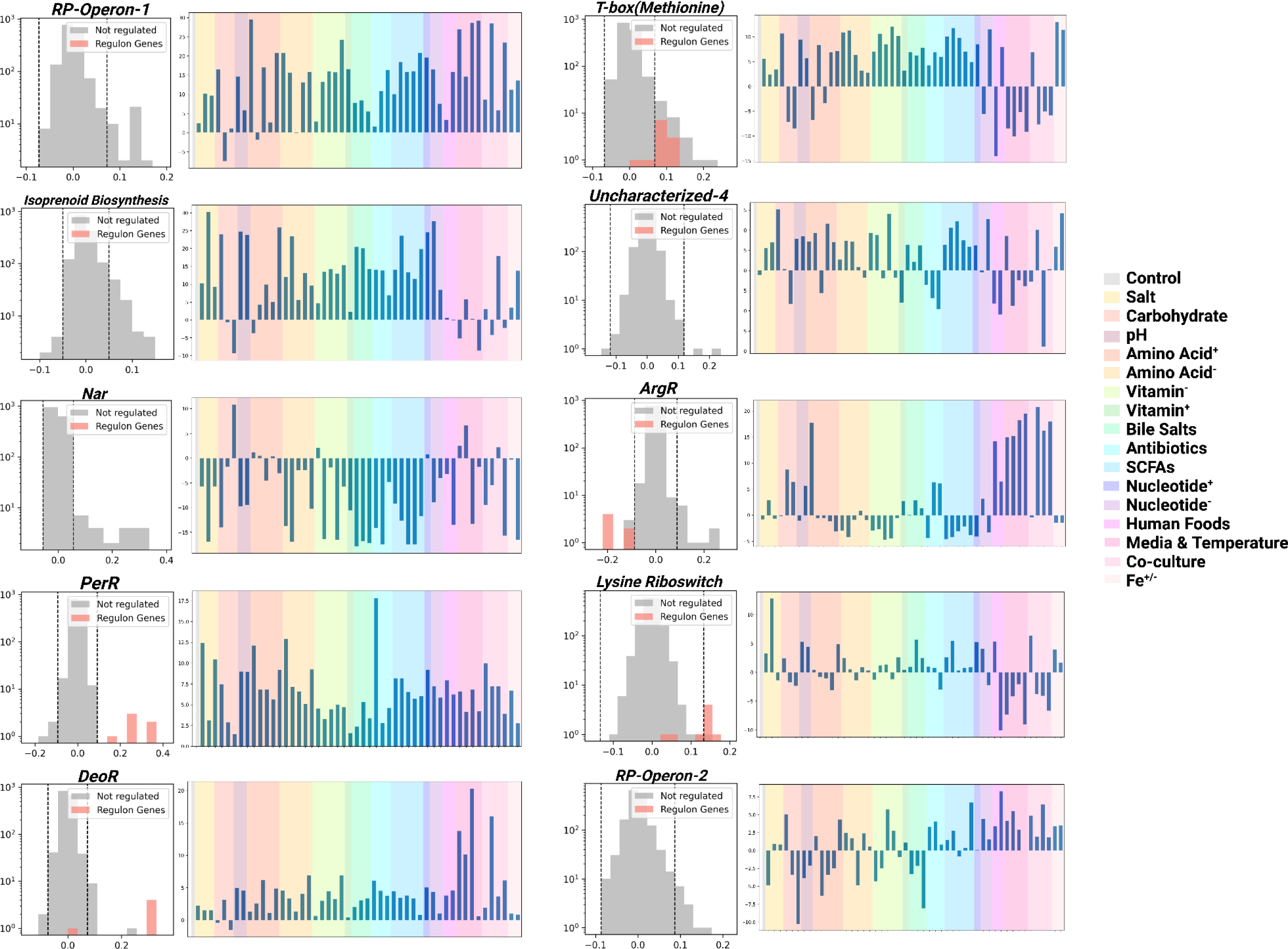
Top-10 iModulons with the highest contribution to the explained variance in the LactoPRECISE compendium. Each subplot presents an iModulon with a histogram detailing gene frequencies across various expression levels. The histogram depicts the distribution of genes within each iModulon (indicated by bars). Adjacent to each histogram is a color-coded bar plot, where colors represent different condition types, and the height of the bars corresponds to the condition’s prominence within the respective iModulon. This visual representation provides insights into the relative influence and condition-specific behavior of each iModulon. For a comprehensive view of the compendium, refer to the **Supplementary** Figure 4, which includes dendrograms and activity plots for the remaining 25 iModulon

iModulon activity, inferred from each respective iModulon across a variety of conditions, is indicative of the biological regulation and signaling that underlies actual activity. Therefore, these activity levels offer valuable insights into the biological function, potentially guiding strain design strategies. It should be noted, however, that the descriptions provided herein are comprehensive but not exhaustive. The application and interpretation of iModulons in the context of microbial foods are wide-ranging, and numerous insights can be gleaned from these results.

In **Figure 2**, we present the top 10 iModulons contributing the most to the explained variance in the LactoPRECISE compendium. Each subplot displays an iModulon and a histogram detailing gene frequencies across different expression levels. The distribution of genes within each iModulon is represented by the histogram bars. Next to each histogram, a color-coded bar plot signifies different condition types, with the bar’s height reflecting the prominence of the condition within the respective iModulon. This visual representation elucidates the relative influence and condition-specific behavior of each iModulon. For a complete view of the compendium, refer to the Supplementary Figures, which include dendrograms and activity plots for the remaining 25 iModulons (**Supplementary Figure 4**).

The iModulons under scrutiny here include RP-Operon-1, Isoprenoid Biosynthesis & Adaptive Response, *Nar*, *PerR*, *DeoR*, T-Box(Methionine), Uncharacterized 4, *ArgR*, Lysine Riboswitch, and RP-operon-2 which all can provide a significant understanding of various aspects of microbial foods.

### Bistable Regulatory Interplay between Fatty Acid Synthesis and Flavin Mononucleotide Production Uncovered by iModulon Analysis

To advance microbial strain design for precision fermentation, achieving control over intricate metabolic networks is crucial. These networks regulate the production of important compounds like vitamins, fatty acids, and natural products in the desired quantities (47, 48, 49, 50). Fatty acid synthesis, specifically, can present unique challenges, such as rate-limiting steps or specific micronutrient requirements for biosynthesis (51, 52). Riboflavin (Vitamin B2) biosynthesis, regulated by particular iModulons, may play a pivotal role in a two-step fatty acid synthesis process. Besides its own nutritional fortification advantages, riboflavin might also influence the production of specific fatty acids, including short-chain and linoleic acids (53, 54,55, 56, 57).

**Figures 3A** and **3B** suggest a coordinated regulatory mechanism by demonstrating concurrent activation or repression of *FabT* and FMN-Riboswitch iModulons under similar conditions. For instance, media containing *E. coli* coculture, starch, pectin, or experiencing thiamine limitation resulted in repressed activities of both iModulons. Conversely, *L. reuteri* grown within media supplemented with glucose at varying pHs or human milk was observed to have increased iModulon activities. The correlation of gene expression levels within these iModulons and their respective activities support this regulatory interconnection.

**Figure 3:**
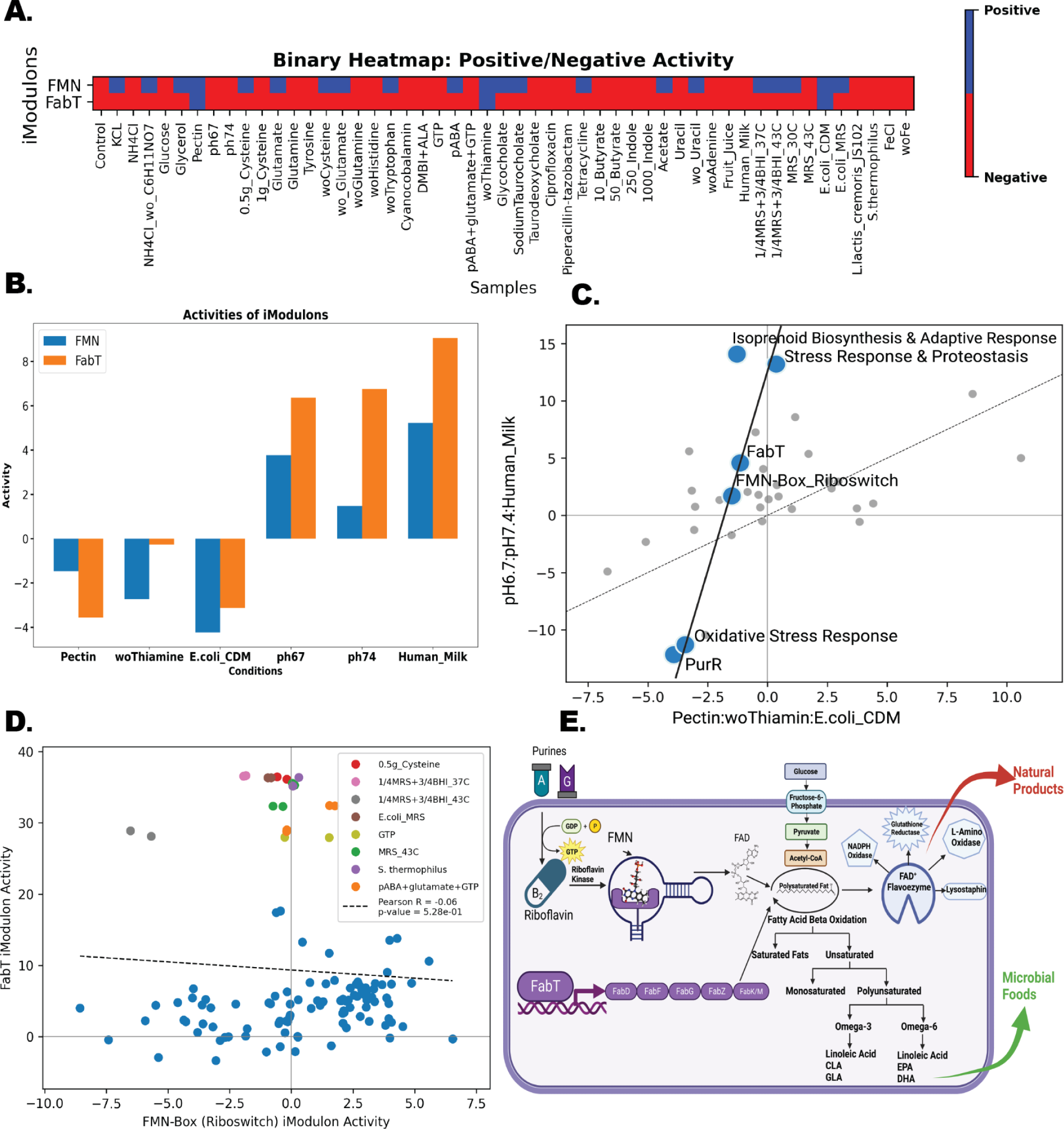
Bistable regulation and interactive metabolic pathways between FabT and FMN Riboswitch iModulons in the production of microbial foods and natural products. (A) Binary heatmap showing the two distinct activities of the FabT and FMN Riboswitch iModulons across various conditions. The color gradient illustrates the relative activity level of the iModulons, with red indicating higher activity and blue indicating negative. Overlapping conditions indicate a transition state between the two iModulons. (B) Bar plot representing conditions that exhibit similar activity between the FMN Riboswitch and FabT iModulons. The conditions are represented on the x-axis, and the corresponding iModulon activity level on the y-axis. This similarity highlights the potential bistability of regulation under these specific conditions. (C) Differential iModulon Activity (DIMA) plot depicting the interactions and shared regulatory trends between various iModulons, including those for stress response and purine production. The distinctive trend line connecting FabT and FMN Riboswitch iModulons reflects their potential bistable regulation. (D) Scatter plot illustrating the iModulon activity of FabT plotted against FMN Riboswitch. The distinctive cluster of conditions at the top suggests a particular set of conditions where both iModulons are simultaneously active, providing evidence for a potential ‘third state’ in the bistable regulation. (E) Schematic diagram depicting the interconnected metabolic pathways involved in the production of microbial foods (mediated by FabT) and natural products (mediated by FMN Riboswitch). This figure highlights the importance of these iModulons in the overall metabolic network and their potential roles in bistable regulation.

This observation is further strengthened by a Differential iModulon Activity (DIMA) plot that illustrates peculiar but similar trajectories for *FabT*, FMN-Riboswitch, and an uncharacterized iModulon under conditions influencing both FabT and FMN-Riboswitch activities (**Figure 3C**). Moreover, **Figure 3D**, a scatter plot illustrating the iModulon activity of FabT plotted against FMN-Riboswitch, reveals a distinctive cluster of conditions at the top (purple outlining). This cluster suggests a particular set of conditions where both iModulons are simultaneously active, providing evidence for a potential ‘third state’ in the bistable regulation, likely due to specific regulatory conditions. Specifically, these highlighted conditions are in mediums that are either supplemented with riboflavin precursors or in anaerobic co-cultures.

Finally, a novel model of cooperative regulation between FMN Riboswitch and FabT iModulons is proposed (**Figure 3E**). Current understanding suggests that these systems are interconnected, with riboflavin supplementation enhancing fatty acid synthesis in probiotics (58). In addition, it is known that riboflavin influences beta oxidation kinetics, a crucial process for cellular ATP production (59). It has been observed in non-prokaryote models that vitamin restrictions or supplements also impacts fatty acid synthesis (60, 61). Thus, we hypothesize an interplay where FMN Riboswitch activation leads to FAD production, which in turn triggers a feedback loop of energy production, fatty acid catabolism, and flavoenzyme formation. This proposed pathway offers insight into the dual role of these iModulons in coordinating the intricate metabolic network necessary for microbial food and natural product production.

### Modulation of Arginine Synthesis by Medium Supplementation: Data-Driven Approach for Enhanced Strain Design

Utilizing insights from microbial transcriptional regulation and assessment of iModulon activity is poised to enhance strain design by providing a nuanced understanding of how different conditions influence microbial phenotypes. In this vein, our study has focused on unraveling the conditions that modulate the activity of the ArgR iModulon, an integral component of arginine synthesis.

The activity of the ArgR iModulon under various conditions was investigated by examining the expression of selected genes: *ArcA*, *ArcC*, *gtfA*, *carA*, *carB*, *argF*, and *argH* (**Figure 4**). By comparing individual gene expression against iModulon activity, we identified conditions that act as activators or repressors of the iModulon. These observations were further strengthened using Pearson’s R correlation, thus providing robust insights into the conditions activating or repressing *ArgR* with high confidence. Co-culturing with Lactic Acid Bacteria (LAB) strains or supplementation with fruit juice consistently activated the *ArgR* iModulon. In contrast, the limitation of various nucleotides or amino acids, exposure to antibiotics, and iron limitation were found to repress the activity of *ArgR*.

**Figure 4:**
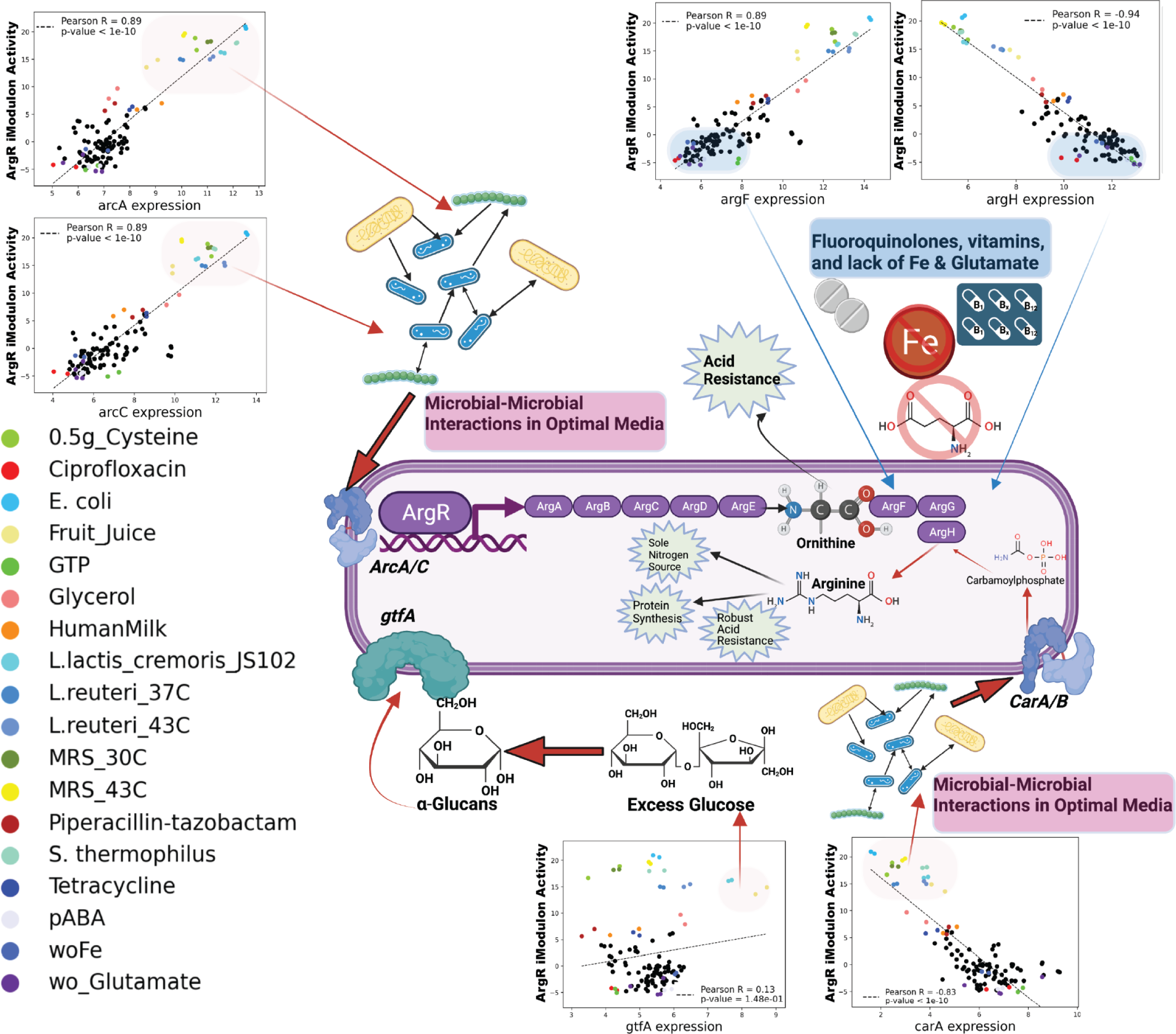
Insight into the context-specific metabolic networks through iModulon analysis, with an emphasis on the *ArgR* iModulon. This figure presents an illustrative diagram of the ArgR iModulon, a key iModulon in the LactoPRECISE compendium. This iModulon comprises various genes, including *arcA*, *arcC*, *argF*, *gtfA*, *argH*, and *carA*. Alongside the schematic, individual gene expression plots corresponding to the activity levels of the *ArgR* iModulon are also displayed. We observe a positive correlation between the high activity of the *ArgR* iModulon and the increased expression of genes such as *arcA*, *arcC*, *argF*, and *gtfA*, suggesting that these genes are co-activated when arginine synthesis is heightened. In contrast, genes *argH* and *carA* display a negative correlation, with high expression when *ArgR* iModulon activity is low, indicating their potential role in regulating the arginine synthesis process. These insights garnered from exploring the complex interplay between iModulon activity and gene expression across different conditions, can be crucial in guiding future strain design applications. It opens up the possibility of fine-tuning these conditions to control *ArgR* iModulon activity and, in turn, arginine synthesis - exemplifying the significant potential of iModulon-based studies in microbial engineering.

Our results have enriched the understanding of the metabolic pathway of *ArgR*. For instance, the presence of various LAB in the culture likely provides excess glutamate, a probable result of positive cross-feeding interactions between the strains (62). Excess glutamate can activate *arcA/C*, facilitating the cellular uptake of glutamate and enabling acid resistance (63). Additionally, excess glycerol in the medium can be taken up by carA or carB, promoting the production of carbamoyl-phosphate, a molecule necessary for *ArgG* function (64).

We also observed that *gtfA* is activated in media supplemented with fruit juice, which expands the existing knowledge that *gtfA* can uptake excess glucose to produce α-glucans (65). α-glucans production by *L. reuteri* may offer advantages such as enhanced acid resistance and amino acid provision for the host and extended food preservation duration (66). In summary, these results demonstrate that iModulon analysis can reveal new insights into cellular metabolism and potentially revolutionize microbial engineering and strain design.

### Decoding *ArgR* Regulation in *L. reuteri* Through Advanced Bitomics

Our study utilized the innovative Bitomics methodology, leveraging comparative structural motifs, to enhance the characterization of the transcriptional unit of *ArgR* in *L. reuteri*, providing valuable insights into its regulatory dynamics (32). This analytical process yielded many insightful findings, advancing our understanding of *ArgR’s* role in this bacterium. The computational methodology was initiated by creating an *L.reuteri* data frame through a novel distance algorithm, which systematically arranged genes into operon clusters. This organization allowed for enhanced comprehension of the spatial arrangement and operational interactions among the genes, thereby setting the stage for downstream analyses. Further insights into the role of *ArgR* were drawn from a gene weight plot for *E. coli* and *L. reuteri ArgR* iModulons. In *E. coli ArgR*, most genes were related to ‘Amino Acid Transport & Metabolism’. However, in *L. reuteri ArgR*, we observed a broader diversity in Clusters of Orthologous Groups (COG) functions and negative gene weights, indicative of a more complex regulatory landscape despite the common *ArgR* motif shared with *E. coli* (**Figure 5A**).

**Figure 5:**
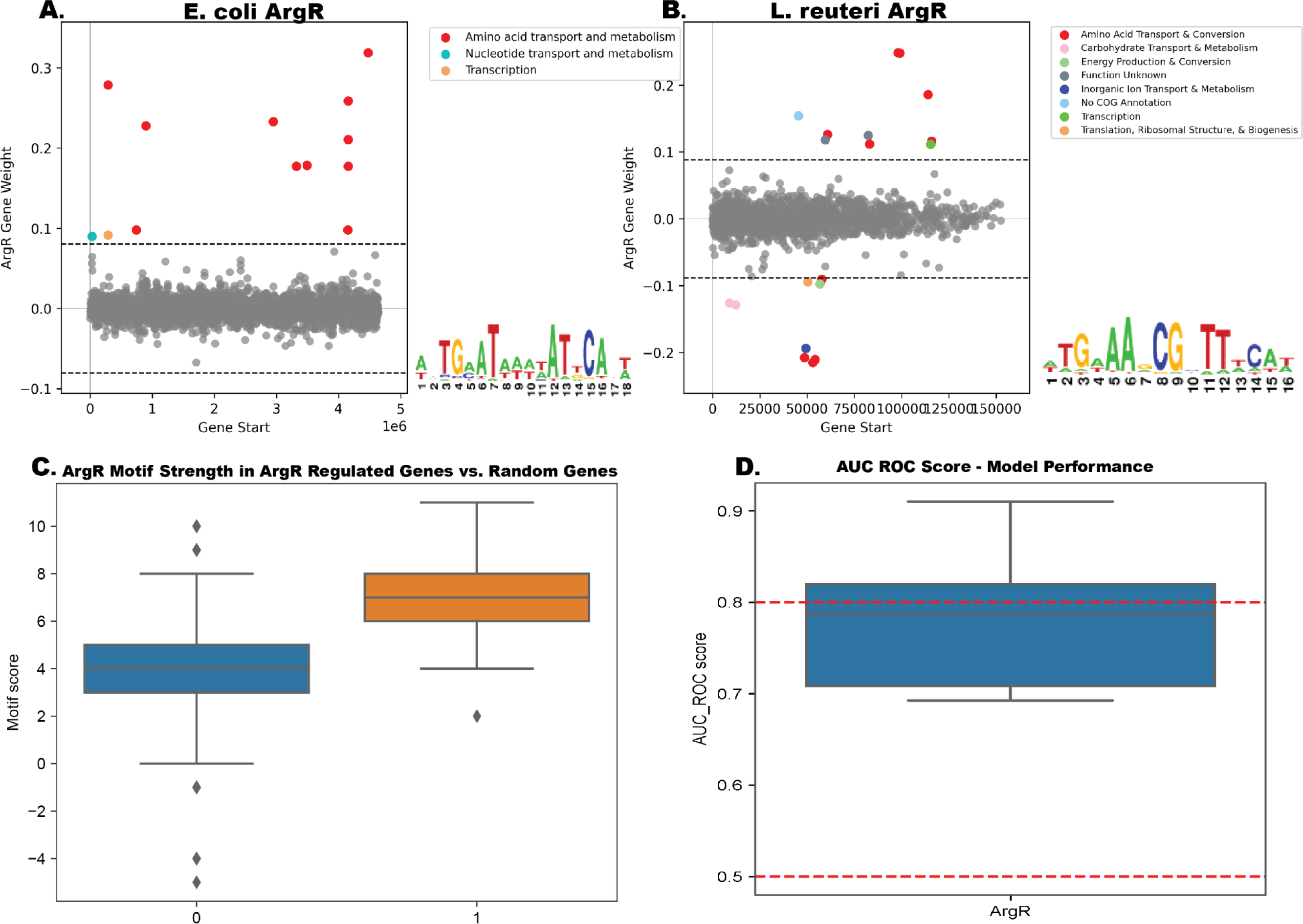
Decoding ArgR Regulation in *L. reuteri* Through Advanced Computational Techniques. (A) Gene weight plot for *E. coli* and *L. reuteri ArgR* iModulons. In *E. coli ArgR*, the majority of genes pertain to ‘Amino Acid Transport & Metabolism.’ Conversely, in *L. reuteri ArgR*, a broader diversity in Clusters of Orthologous Groups (COG) functions and negative gene weights is observed, suggesting a more complex regulatory scenario despite the similar ArgR motif between the two organisms. (B) Analysis of *ArgR* motif strength in *ArgR* regulated genes versus random genes. This plot presents a comparison of ArgR motif prevalence and strength among genes regulated by *ArgR* and a random selection of genes. (C) Area Under the Receiver Operating Characteristic (AUC-ROC) score assessing the performance of the computational model used in this study. A high AUC-ROC score reflects the model’s capacity to classify genes correctly under the ArgR regulation based on motif strength. The combined analysis presented in this figure illustrates the complexity of *ArgR* regulation in *L. reuteri* and points towards potential differences in regulatory strategies compared to *E. coli,* a thoroughly studied model organism

A Position-Specific Scoring Matrix (PSSM) was also generated to identify the distribution of nucleotide frequencies at each position within the ArgR binding sites. Using the log-odds scores derived from the PSSM, we could identify statistically significant motifs within the gene sequences. With these motif scores, a logistic regression model, bolstered by elastic net regularization, was trained. The performance of this model was exceptional, with an Area Under the Receiver Operating Characteristic Curve (AUC-ROC) score nearing 0.8, attesting to the model’s robust predictive capacity for ArgR regulation (**Figure 5B**). The class imbalance was successfully mitigated using a resampling technique, contributing to a balanced training set that further strengthens the model’s robustness.

A marked distinction was observed in the motif scores of genes residing within and outside the *ArgR* motif. Genes within the *ArgR* motif presented higher motif scores, thereby confirming our model’s predictive accuracy regarding *ArgR* regulation in *L. reuteri*. A comparison of *ArgR* motif prevalence and strength between genes regulated by *ArgR* and a random gene selection further underscored this finding (**Figure 5C**). In summary, we successfully employed advanced Bitomics to gain unprecedented insights into the *ArgR* regulatory dynamics in *L. reuteri*. By leveraging machine learning and statistical modeling techniques, we have opened avenues for future research into the TRNs of other industrially significant microbes.

### Interplay between iModulon Activity and Nitrogen Metabolism in Secondary Metabolite Production

Our investigations into the complex interactions between the Type III Polyketide Synthase (T3PKS) gene cluster and iModulon activity in *L. reuteri* revealed novel insights into nitrogen metabolism and isoprenoid biosynthesis. As depicted in **Figure 6A**, the T3PKS gene cluster, predicted by antiSMASH 7.0 (43), is composed of four active genes involved in polyketide synthesis, a group of secondary metabolites with considerable biological diversity and potential therapeutic applications (67).

**Figure 6:**
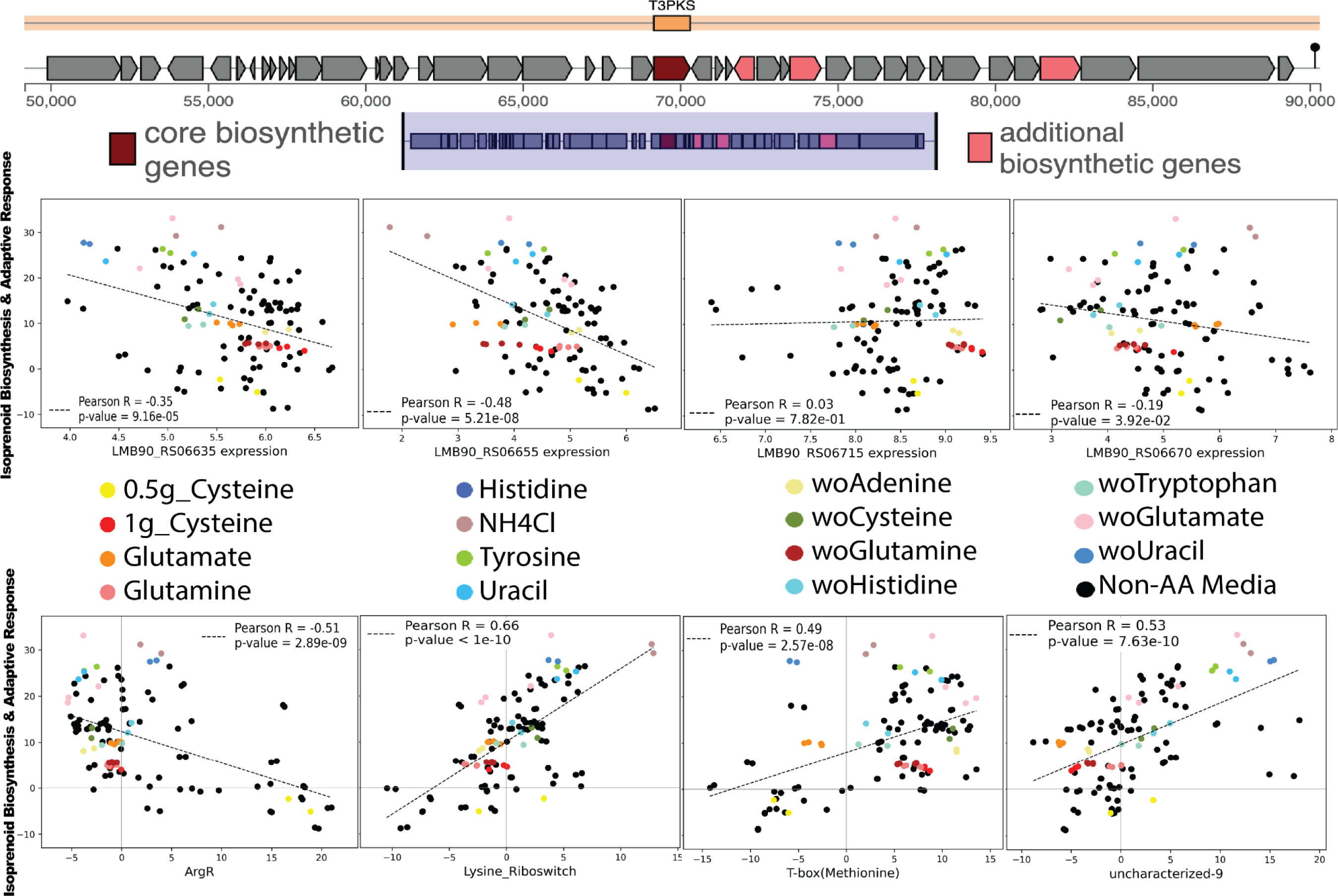
Unraveling the interplay between the Type III Polyketide Synthase (T3PKS) gene cluster and iModulon activity in *L. reuteri,* with an emphasis on nitrogen metabolism and isoprenoid biosynthesis. (A) showcases the predicted biosynthetic gene clusters (BGCs) from the *L. reuteri* genome, as predicted by antiSMASH 7.0. The focus is on the T3PKS gene cluster, which comprises one core biosynthetic gene and three additional biosynthetic genes found to be active in our compendium. T3PKS enzymes are involved in synthesizing polyketides, a large family of secondary metabolites with diverse biological activities and therapeutic potential. (B) we present the gene expression of these four biosynthetic genes plotted against the ‘Isoprenoid Biosynthesis and Adaptive Response’ iModulon. This iModulon encapsulates genes involved in the production of isoprenoids - critical components of cell membranes - and genes associated with stress response and adaptation. The plots underscore the strong activation of this iModulon under conditions related to nitrogen metabolisms, such as the removal of glutamate or uracil and supplementation with NH4Cl. (C) further explores this iModulon activity in relation to three well-characterized iModulons (*ArgR*, Lysine Riboswitch, and T-box (Methionine)), all of which are known to regulate amino acid metabolism, as well as the ‘uncharacterized-9’ iModulon, encompassing genes involved in ammonia transport. Strong correlations emerge between the ‘Isoprenoid Biosynthesis and Adaptive Response’ iModulon and these four iModulons, reinforcing the role of nitrogen metabolism in regulating secondary metabolite production. This figure illuminates potential strategies for engineering *L. reuteri* to optimize the production of secondary metabolites by modulating conditions that highly activate key iModulons.

Interestingly, the ‘Isoprenoid Biosynthesis and Adaptive Response’ iModulon demonstrated strong activation when correlated with the expression of these four biosynthetic genes (**Figure 6B**). This iModulon comprises genes vital for isoprenoid production - essential constituents of cell membranes - and for the organism’s adaptive stress response. Under specific conditions tied to nitrogen metabolism, including glutamate or uracil depletion and NH4Cl supplementation, this iModulon was notably active.

**Figure 6C** further delves into the interconnections between this iModulon activity and four well-characterized iModulons. These iModulons - ArgR, Lysine Riboswitch, and T-box (Methionine) - are known regulators of amino acid metabolism. Additionally, the ‘uncharacterized-9’ iModulon, consisting of genes associated with ammonia transport, was explored. Robust correlations emerged between the ‘Isoprenoid Biosynthesis and Adaptive Response’ iModulon and these four iModulons, underscoring the significant role nitrogen metabolism plays in modulating secondary metabolite production (68). Interestingly, hierarchical clustering of the top-five most significant iModulon clusters in LactoPRECISE highlights that Cluster 2 contains the iModulons ‘Isoprenoid Biosynthesis and Adaptive Response’, ‘Lysine Riboswitch’, and ‘Uncharacterized-9’; these results support the notion of co-regulation of these iModulons and a shared response across the compendium (**Supplementary Figure 3)**. These findings offer crucial strategies for the potential genetic engineering of *L. reuteri* to maximize secondary metabolite production, achieved by adjusting conditions to activate key iModulons.

## V. Discussion/Conclusion

In this study, we have successfully managed to delineate and reconstruct the TRN of *L. reuteri*. This was achieved by identifying 35 distinct iModulons using ICA applied to a comprehensive set of 117 high-quality RNA-seq samples. A closer examination led to the discovery of 13 Regulatory iModulons with significant links with recognized Transcription Factors (TFs). These findings not only validate but also extend upon previously known TF-gene interactions, bridging the gap in our understanding of potential co-regulators and regulatory targets through the systemic analysis of iModulon gene memberships.

Our approach shed light on 11 unique clusters of genes, termed Functional iModulons. The importance of these iModulons is underscored by their co-expression during environmental changes, making them promising focal points for future research geared towards unearthing yet unidentified regulators. In contrast with traditional methodologies that isolate a few experimental conditions for analysis, ICA enables simultaneous evaluation of varied gene expression profiles. This helps to identify clusters of genes most likely regulated by specific TFs in a highly efficient manner.

The iModulon-centric method employed in our study, complemented by others within iModulonDB, signifies that examining iModulons provides a powerful, insightful perspective to observe significant shifts in microbial gene expression. This comprehensive, systems-level perspective offers a more holistic understanding of global changes in gene expression profiles, an aspect often overlooked by traditional RNA-seq studies.

The TRNs and iModulons generated from specific gene expression profiles of non-model organisms like *L. reuteri* serve as invaluable tools to characterize organisms with limited gene annotation and regulatory network information. In this study, we successfully optimized the dimensionality of iModulons to strike a balance between the over- and under-decomposition of our gene expression compendium. This optimization enriches the interpretability of newly sequenced transcriptomes and elucidates dynamic shifts in transcriptomes across different conditions. These conditions often mirror those found in human-associated and commercially significant bioprocesses, further increasing the relevance of our findings.

By studying the activities of iModulons, we were able to generate new hypotheses, particularly about how different conditions influence the expression of genes regulated by the *ArgR* iModulon, vital for arginine production. However, our research is not devoid of limitations. The utility of iModulon characterization can be significantly enhanced by applying ICA to more diverse and larger RNA-seq sample sets. Future endeavors in this direction will not only improve our understanding of modules that comprise *L. reuteri*’s TRN, but also offer valuable insights into other probiotic strains.

In conclusion, our study represents a pioneering global gene expression analysis across a broad spectrum of conditions for *L. reuteri*. The insights derived from the 35 identified iModulons form the bedrock of a more advanced understanding of the biological intricacies of *L. reuteri* and open doors for practical applications in strain design. The findings of this study underscore the immense potential of deploying systems biology tools to elucidate the complex landscape of microbial gene expression and regulation. We stand on the cusp of a future where these insights will be crucial in tailoring probiotics for specific health outcomes and manipulating microbial consortia in food systems, truly transforming the future of microbial foods.

## DATA/CODE AVAILABILITY

All in-house generated sequences were deposited in the NCBI-Sequence Read Archive database (PRJNA989027). The accession number of the deposited reads is provided in **Supplementary Table 1**. While the X, M, and A matrices in addition to TRN regulator file, gene ontology, iModulon Table Files, and code to produce these are all available on GitHub (https://github.com/PoeticPremium6/LactoPRECISE). A DOI with the code used in this manuscript can also be found on Zendo (https://zenodo.org/record/8108422). Each gene and iModulon have interactive, searchable dashboards on iModulonDB.org https://imodulondb.org/dataset.html?organism=l_reuteri&dataset=lactoprecise, and data can also be downloaded from there.

## Acknowledgements

We are grateful to Omkar Satyavan Mohite for the informative discussion on biosynthetic gene cluster analysis. We thank Marc Abrams for reviewing the manuscript and providing constructive suggestions. We are also thankful to Rebecca Höög for improving the quality of all figures.

