## Supplementary Figures for "Reconstructing the Transcriptional Regulatory Network of Probiotic *L. reuteri* is Enabled by Transcriptomics and Machine Learning"

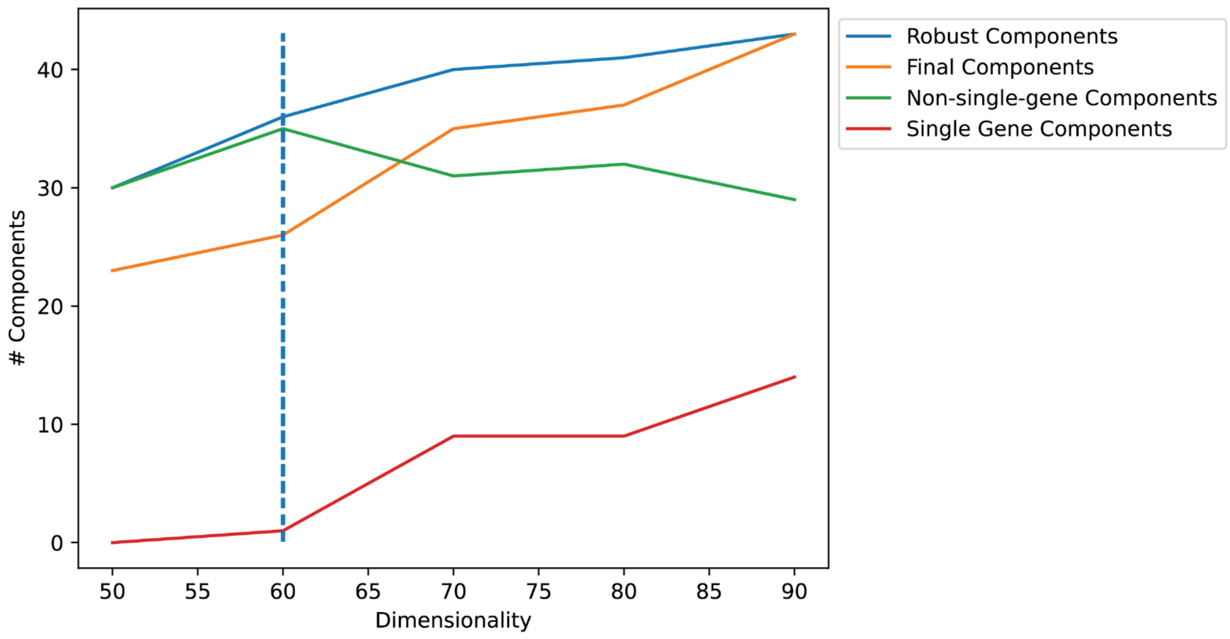

Robust Components
Final Components
Non-single-gene Components
Single Gene Components
### Components
Dimensionality
0
10
20
30
40
50
55
60
65
70
75
80
85
90


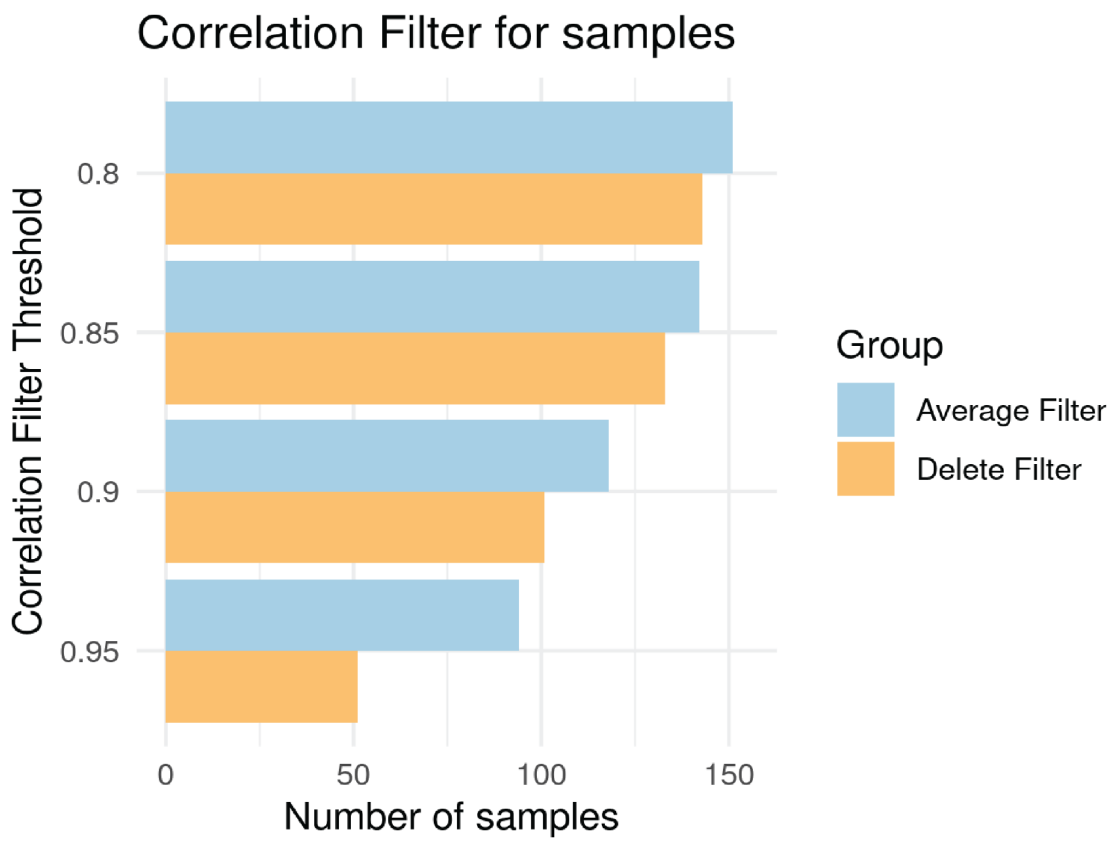

Correlation Filter for samples
Correlation Filter Threshold
0.8
0.85
0.9
0.95
0
50
100
150
Number of samples
Group
Average Filter
Delete Filter

**Supplementary Figure S1:Identification and evaluation of optimal dimensionality for independent components in the iModulon analysis of L. reuteri.** A) The Python script 'get_dimension.py' was used to perform an iterative process of component extraction and evaluation in order to determine the dimensionality that results in the most robust and meaningful iModulon components. The figure plots the number of components identified against their respective dimensionalities. Four types of components are illustrated: robust components, final components, non-single-gene components, and single-gene components. Robust components represent all components found in the analysis, while final components are those that meet a certain similarity threshold compared to the highest dimension. Non-single-gene components and single-gene components distinguish components based on whether they contain multiple genes or are driven by a single gene, respectively. A vertical dashed line indicates the optimal dimensionality that maximizes the number of final components while maintaining the most multi-gene components. This dimensionality value provides a balance between the complexity of the model (number of dimensions) and the interpretability of the components (number of multi-gene components). The results from this analysis, particularly the matrices of components (M.csv) and their activities (A.csv) at this optimal dimensionality, serve as the basis for subsequent iModulon characterization and interpretation.
B) This figure illustrates the results of a correlation-based filtration and averaging procedure performed on gene expression data. Gene expression data were grouped according to predefined conditions, and the pairwise Pearson correlation between samples within each group was computed. For any pair of samples within a group exhibiting a correlation coefficient below a specified threshold (values used: 0.8, 0.85, 0.9, 0.95), all samples in the group were averaged to create a representative expression profile for that group. The output of this procedure is a filtered expression matrix where highly dissimilar samples have been amalgamated into single profiles. Each subfigure corresponds to a different correlation threshold used in the filtration process.

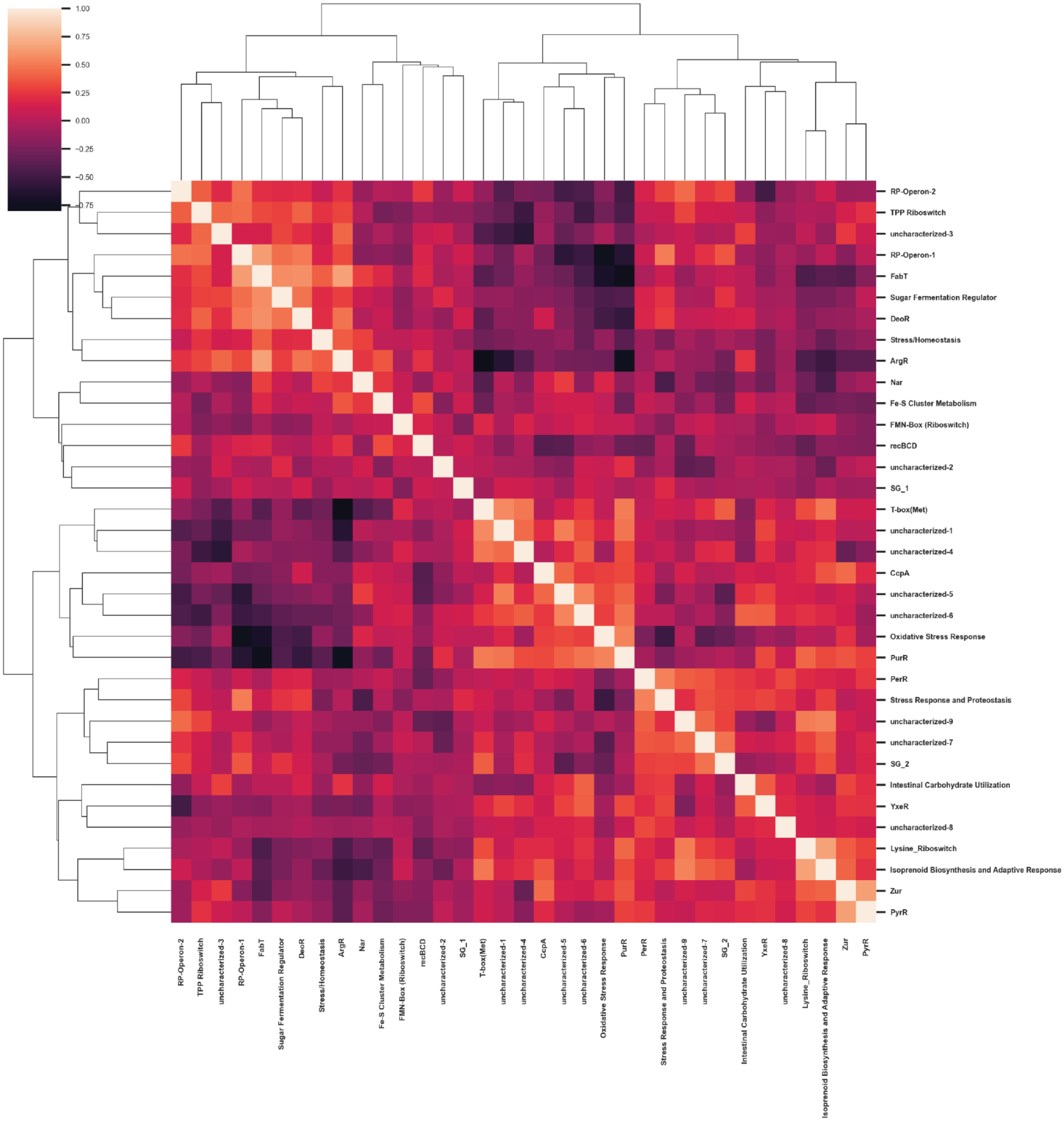

1.00
0.75
0.50
0.25
0.00
−0.25
−0.50
−0.75
RP-Operon-2
TPP Riboswitch
uncharacterized-3
RP-Operon-1
FabT
Sugar Fermentation Regulator
DeoR
Stress/Homeostasis
ArgR
Nar
Fe-S Cluster Metabolism
FMN-Box (Riboswitch)
recBCD
uncharacterized-2
SG_1
T-box(Met)
uncharacterized-1
uncharacterized-4
CcpA
uncharacterized-5
uncharacterized-6
Oxidative Stress Response
PurR
PerR
Stress Response and Proteostasis
uncharacterized-9
uncharacterized-7
SG_2
Intestinal Carbohydrate Utilization
YxeR
uncharacterized-8
Lysine_Riboswitch
Isoprenoid Biosynthesis and Adaptive Response
Zur
PyrR

**Supplementary Figure S2: IModulon Correlation Heatmap.** The figure displays a cluster map representing the correlation among selected iModulons in L. reuteri. This heatmap utilizes hierarchical clustering to group iModulons based on similarities in their gene expression patterns across various conditions. The dendrogram and bold labels emphasize strong correlations, providing insight into potential shared or complementary roles in cellular responses.

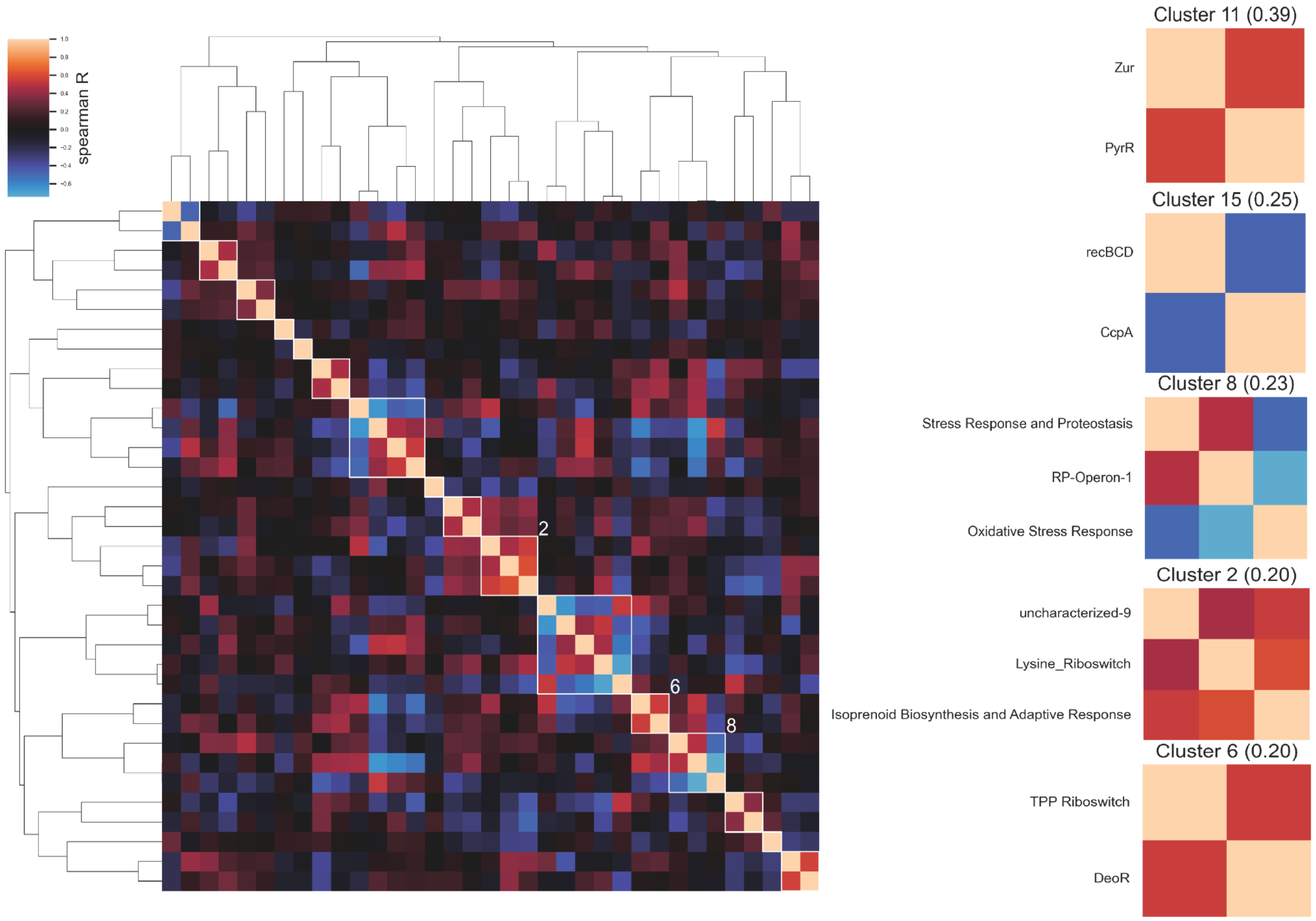


**Supplementary Figure S3: Hierarchical clustering of the five most significant iModulon clusters in L. reuteri.** Each sub-figure within the set represents a distinct iModulon cluster and its corresponding activity profile across various conditions. These visuals highlight the clusters that exhibit the most substantial and consistent patterns, providing insight into the potential co-regulation of iModulons and their collective impact on L. reuteri's cellular responses.

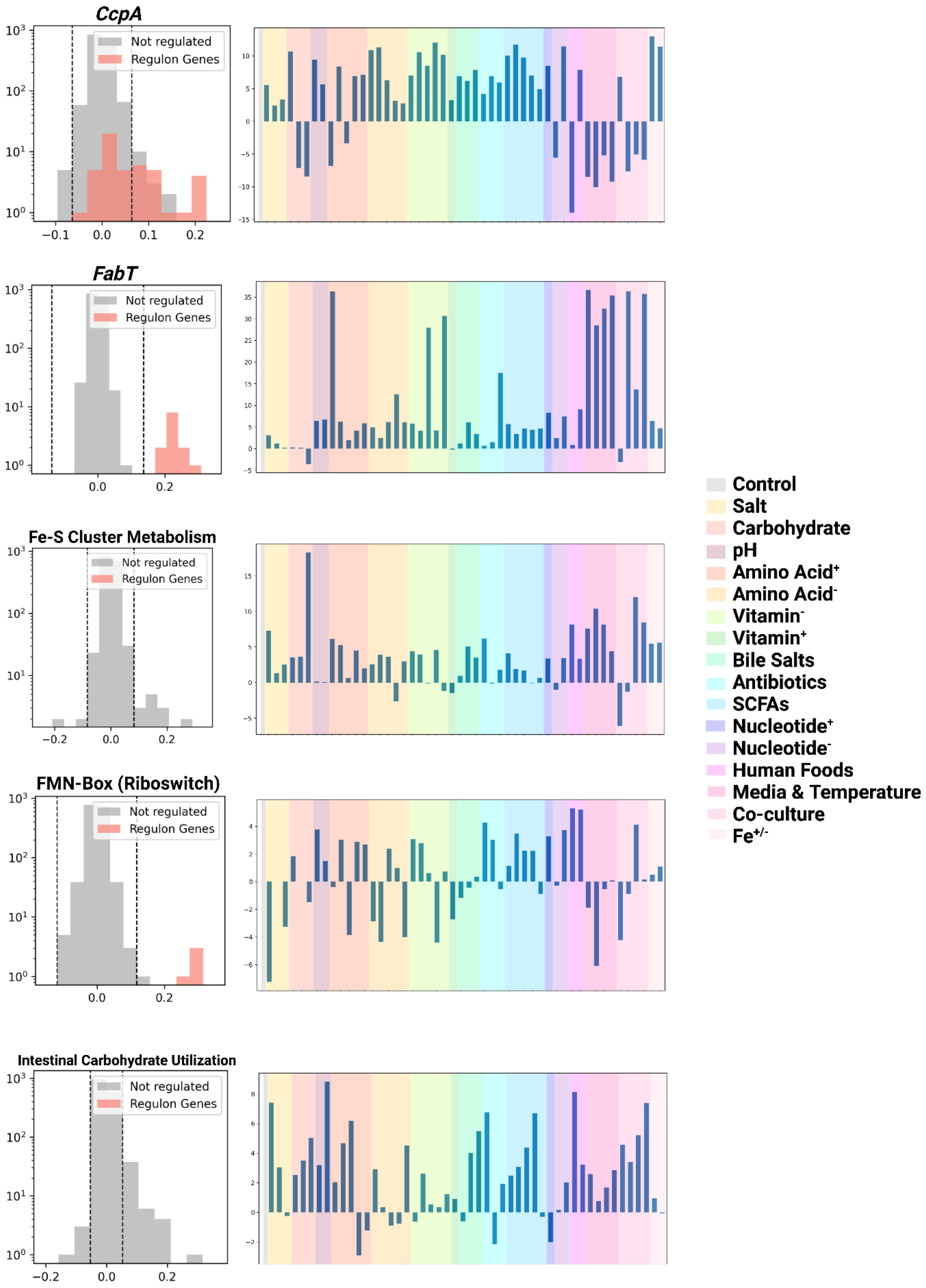

CcpA
FabT
Fe-S Cluster Metabolism
FMN-Box (Riboswitch)
Intestinal Carbohydrate Utilization
Not regulated
Regulon Genes
Control
Salt
Carbohydrate
pH
Amino Acid+
Amino Acid-
Vitamin-
Vitamin+
Bile Salts
Antibiotics
SCFAs
Nucleotide+
Nucleotide-
Human Foods
Media & Temperature
Co-culture
Fe+/-

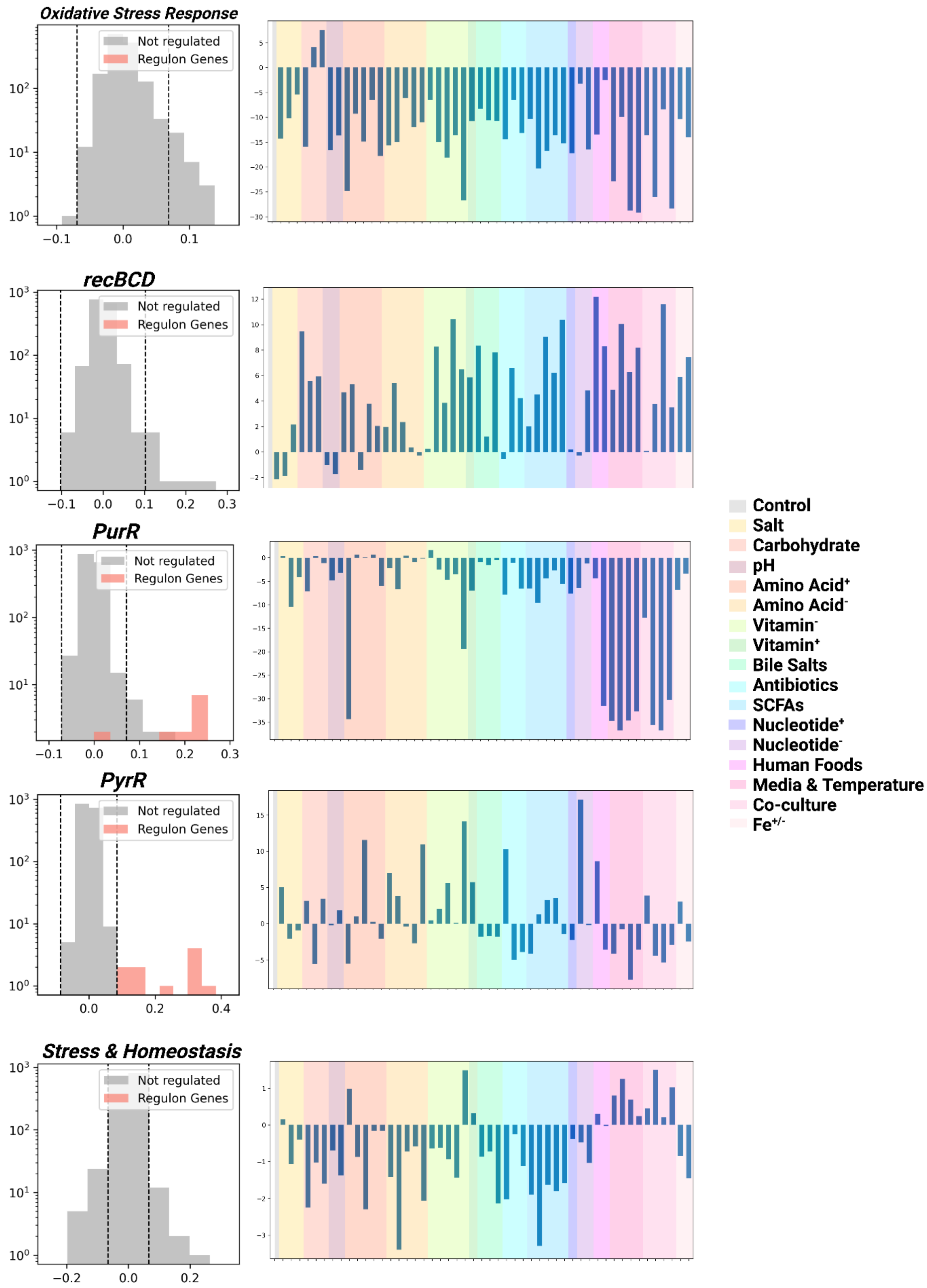
Oxidative Stress Response
recBCD
PurR
PyrR
Stress & Homeostasis
Not regulated
Regulon Genes
Control
Salt
Carbohydrate
pH
Amino Acid+
Amino Acid-
Vitamin-
Vitamin+
Bile Salts
Antibiotics
SCFAs
Nucleotide+
Nucleotide-
Human Foods
Media & Temperature
Co-culture
Fe+/-

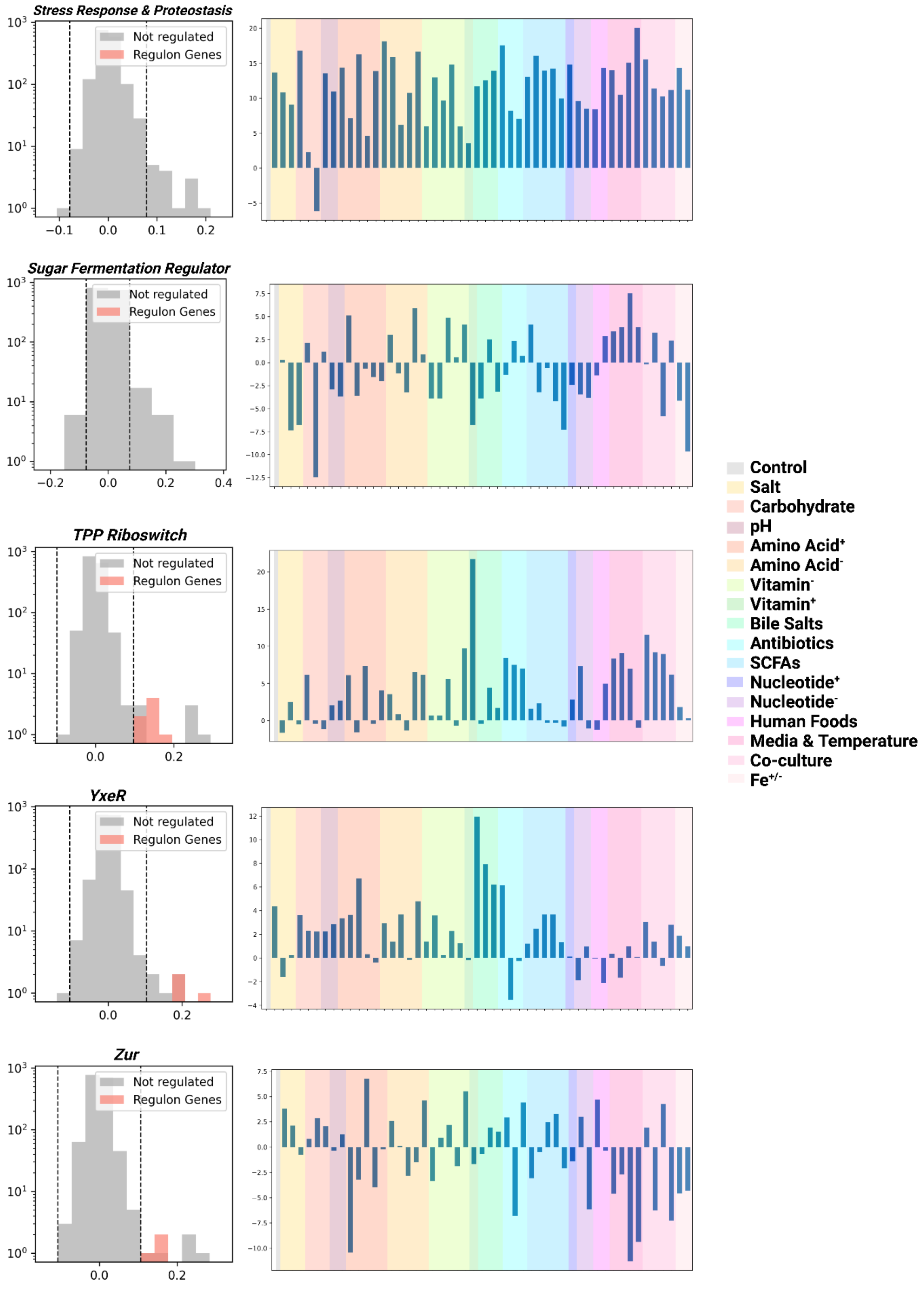

Stress Response & Proteostasis
Sugar Fermentation Regulator
TPP Riboswitch
YxeR
Zur
Not regulated
Regulon Genes
Control
Salt
Carbohydrate
pH
Amino Acid+
Amino Acid-
Vitamin-
Vitamin+
Bile Salts
Antibiotics
SCFAs
Nucleotide+
Nucleotide-
Human Foods
Media & Temperature
Co-culture
Fe+/-

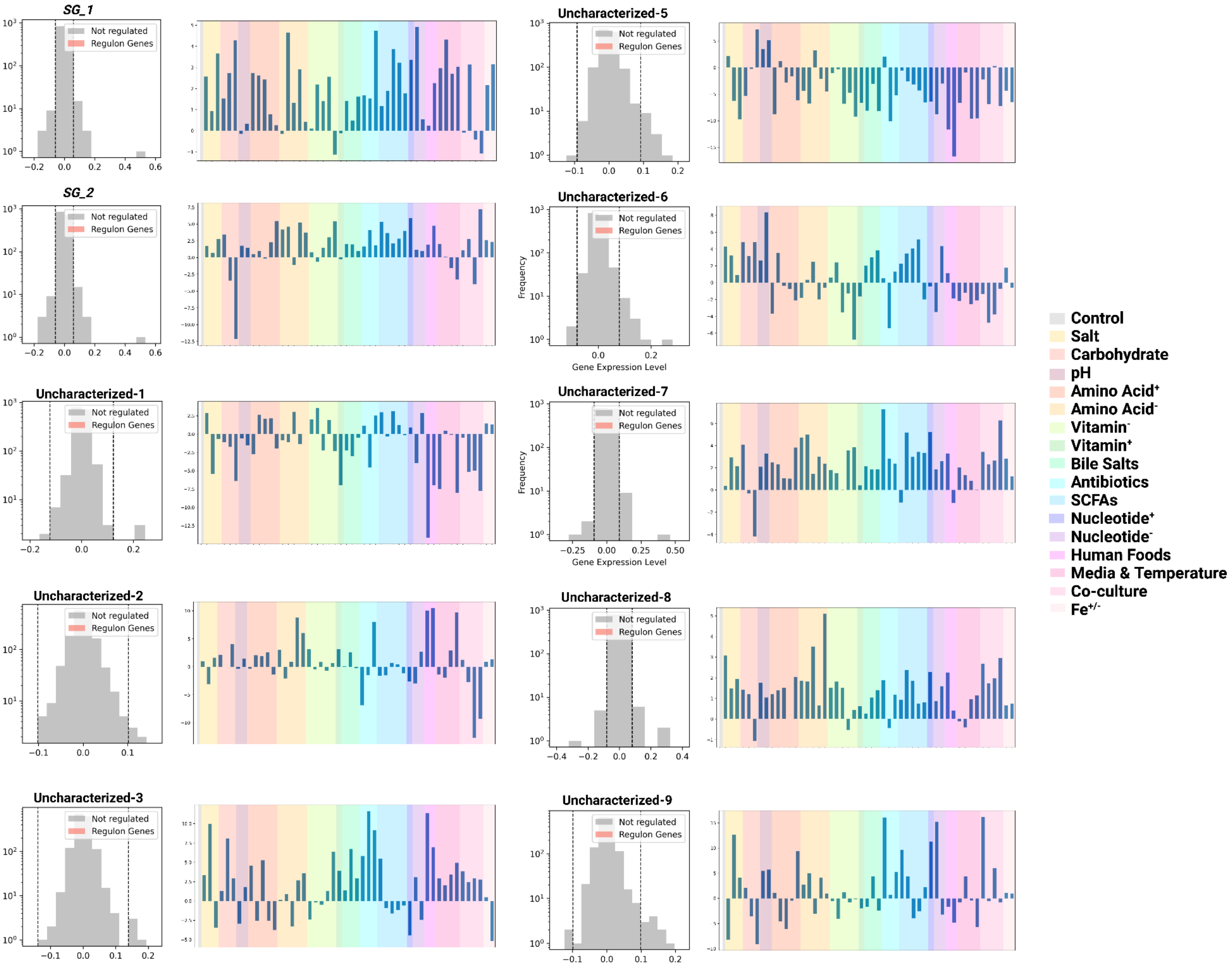

SG_1
SG_2
Uncharacterized-1
Uncharacterized-2
Uncharacterized-3
Uncharacterized-5
Uncharacterized-6
Uncharacterized-7
Uncharacterized-8
Uncharacterized-9
Not regulated
Regulon Genes
Frequency
Gene Expression Level
Control
Salt
Carbohydrate
pH
Amino Acid+
Amino Acid-
Vitamin-
Vitamin+
Bile Salts
Antibiotics
SCFAs
Nucleotide+
Nucleotide-
Human Foods
Media & Temperature
Co-culture
Fe+/-

**Supplementary Figure S4**: Remaining 25 iModulons that were identified in the LatoPRECISE compendium
